# From forests to farming: identification of photosynthetic limitations in breadfruit across diverse environments

**DOI:** 10.1101/2024.07.08.602079

**Authors:** Graham J. Dow, Noa Kekuewa Lincoln, Dolly Autufuga, Robert Paull

## Abstract

Breadfruit (*Artocarpus altilis*) is a prolific tropical tree producing highly nutritious and voluminous carbohydrate-rich fruits. Already recognized as an underutilized crop, breadfruit could ameliorate food insecurity and protect against climate-related productivity shocks in undernourished equatorial regions. However, a lack of fundamental knowledge impedes widespread agricultural adoption, from modern agroforestry to plantation schemes. Here, we used a multi-environment breadfruit variety trial across the Hawaiian Islands to determine photosynthetic limitations, understand the role of site conditions or varietal features, and define their contributions to agronomic efficiency. Photosynthetic rates were dependent on location and variety, and strongly correlated with fruit yield (r^2^=0.80, p<0.001). Photochemistry was suitable to full-sunlight conditions, with a saturation point of 1545 PAR, *V*_cmax_ of 151 μmol m^-2^s^-1^, and *J*_max_ of 128 μmol m^-2^s^-1^, which are high-end compared to other tropical and temperate tree crops. However, limitations on CO_2_ assimilation were imposed by stomatal characteristics, including stomatal density (p<0.05) and diurnal oscillations of stomatal conductance (>50% reductions from daily maxima). These constraints on CO_2_ diffusion are likely to limit maximum productivity more than photochemistry. Our results comprise the first comprehensive analysis of breadfruit photosynthesis, successfully links ecophysiology with fruit yield, and identifies vital traits for future research and management optimization.

## Introduction

Neglected and underutilized species (NUS) have tremendous potential to contribute to food security, sovereignty, and sustainability, although substantial barriers still exist to realize that potential (Padulosi et al., 2002, 2021). Arguably, one reason is that many NUS have been selected to operate in cropping systems that have been displaced by industrial monocultures (Lincoln et al., 2023). Therefore, NUS can reflect traits that specifically excel within traditional ecologically adapted cropping systems, either because they were selected to perform in such systems or because they are ‘semi-domesticated’ and still reflect traits related to their wild habitat (Harlan, 1992). Some applications of NUS seek to employ them in conventional (i.e., monocropped, high-input) systems when this may not be the agricultural form to which they are best adapted.

In particular, many NUS crops were used in various agroforestry systems. Just as forests represent a particular point on a spectrum of ecosystem types (grasslands, shrublands, etc.), agroforestry systems represent a point on a spectrum of agroecological systems, from intensive monocropping of annuals to diversified, multistrata perennial systems. In both natural and agricultural ecosystems, systems differ in the distribution of many environmental parameters, such as light intensity, humidity, temperature, and available soil nutrients. Plants within each system adapt to the associated parameters depending on their locational situation (i.e., canopy or understory, proximity to neighbors, etc.) and strategy (i.e., pioneer species, slow growth, etc.) through physiological parameters such as light saturation, stomatal function, and nutrient use efficiency. NUS offer an opportunity to consider the shift in physiological traits in selection for various agroecosystems.

One such NUS is breadfruit (*Artocarpus altilis* (Parkinson) Fosberg), a broadleaf tropical tree in the Moraceae family that produces large fruits rich in complex carbohydrates that are used as a staple crop on many Pacific islands (Ragone, 1997). The trees were traditionally grown in various cropping systems, ranging from closed canopy food forests to part of diversified agroforestry systems, to individual home trees (Meilleur et al., 2024; Quintus et al., 2019). Breadfruit evolved from its wild ancestor breadnut (*Artocarpus camansi*), a forest-dwelling species native to New Guinea and perhaps surrounding islands (Zerega et al., 2006). The selection of cultivars was accelerated by human-mediated island hopping, with the repeated selection and isolation of subsets creating a rapid evolution of breadfruit domestication, as seen in a clear spectrum of seediness (Ragone, 1997) and mycorrhizal associations (Xing et al., 2012) as one radiates from Papua New Guinea. Currently, breadfruit has been recognized as a priority crop for addressing global hunger and food security (Lucas and Ragone, 2012) and is increasingly grown in monocultured orchards (Lincoln et al., 2018). However, such a transition may require adaptation or optimization of traits for high-light environments, or conversely, the application of more appropriate cropping systems.

Breadfruit has traditionally been used as a staple in the Pacific Islands for more than 3,000 years (Ragone, 1997). The plant is well adapted to many tropical climates and does especially well in the wet tropics, where many other staple crops, especially grains, do not. Large, rapidly growing trees have a unique leaf structure, producing large leaves up to a meter in length, typically with deep lobes that in extreme cases can extend to the mid-rib. Trees can produce significant amounts of starchy fruits, with annual yields of 300-600 lbs per tree common and up to 2,000 lbs per tree reported (Meilleur et al., 2024).

Due to its high productivity and contributions to human nutrition, including a high complex carbohydrate content and complete amino acid profile (Liu et al. 2015; Needham et al. 2020), breadfruit has been identified as one of the 35 priority underutilized crops for hunger and climate adaptation (FAO, 2009). The breadfruit genome is enriched with genes for photosynthesis, starch, and sucrose metabolism compared to closely related species (Sahu et al., 2020), providing a tremendous genetic platform for crop improvement. As more than 80% of the undernourished people live in tropical and subtropical regions and food insecurity is only expected to increase due to changing economic and climatic conditions (Bell et al., 2016), breadfruit has the potential to help alleviate hunger, increase food security, and improve human nutrition (Jones et al., 2013, Meilleur et al., 2024, Liu et al., 2015, 2020).

Although the historical significance and agricultural potential of breadfruit is obvious, characterization of basic plant physiology, trait variation, and environmental responses is relatively lacking, which is true for many NUS. Some attempts have been made to fill significant knowledge gaps, for example, net photosynthesis, stomatal conductance, and the transpiration rate of breadfruit are less affected by salt stress than its relative jackfruit (Su et al., 2019). Furthermore, seasonal changes in breadfruit leaf chlorophyll content and photosynthesis are influenced by sensitivity to chilling, with photosynthetic rates ranging from 10.7 to 17.1 µmol CO_2_ m⁻² s⁻¹ (Cao et al., 2006). Overcoming limited data on breadfruit photosynthesis under field conditions and the influence of the environment on carbon assimilation, water use, and fruit development are crucial for the expansion of breadfruit production areas and agronomic resiliency.

To understand the limitations on breadfruit productivity in agricultural systems, we evaluated the leaf physiology of five breadfruit varieties and the wild ancestor (breadnut) in full-sunlight orchard schemes across multiple environments. Our objective is to characterize photosynthetic assimilation in breadfruit varieties to provide information on tree growth and fruit yield, as well as carbon sequestration potential and water demands. Our data capture multiple aspects of carbon assimilation, including biochemical parameters and saturation points, diffusion limitations, canopy effects, site and variety differences, and diurnal patterns. This is, to our knowledge, the first comprehensive assessment of carbon assimilation data for breadfruit, or any *Artocarpus* species. Ultimately, this research identifies traits that could improve the intrinsic physiology of breadfruit trees and management approaches that would benefit agricultural productivity.

## Materials and Methods

### Study sites

This study took advantage of an established variety trial network in Hawai‘i (Lincoln et al. 2019a). Each location consists of 30 trees at 10m (30ft) spacing, consisting of five plantings each of five breadfruit varieties (cvs. Ma’afala, Fiti, Otea, Pua’a and ‘Ulu Maoli) and the wild ancestor *Artocarpus camansi* (breadnut). We used six sites distributed across soils and environments in the Hawaiian archipelago (Figure 1; Table 1). An additional site located at the Waimanalo Research Station (21.3355° N, 157.7148 ° W) on Oahu with mature plantings of the Ma’afala variety established in 2014 was used to evaluate diurnal patterns and specific response curves, but was not included in the comparisons between environments.

**Figure 1:**
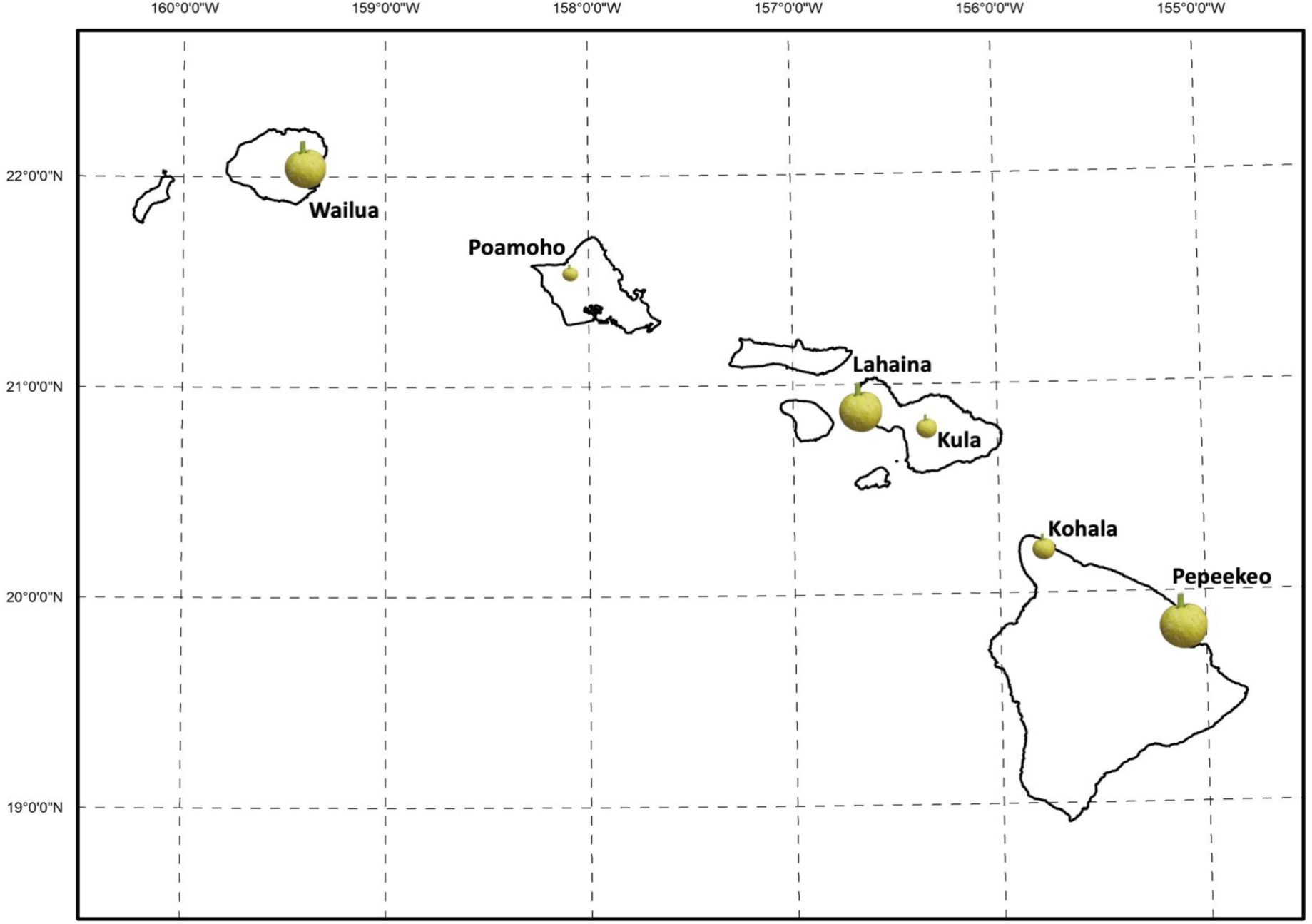
Locations of the variety trial sites used in this study and relative fruit yields. The size of the fruit is proportional to the average yield for each site transformed by square root.

**Table 1:** Site locations used in this study and associated environmental characteristics.

| **Site** | **Elevation**<br>**(m)** | **Rainfall**<br>**(mm/y)** | **ET**<br>**(mm/y)** | **Net Rad**<br>**(W/m<sup>2</sup>)** | **Cloud**<br>**Freq (%)** | **Temp**<br>**(°C)** | **VPD**<br>**(Pa)** | **RH**<br>**(%)** |
| --- | --- | --- | --- | --- | --- | --- | --- | --- |
| Kohala | 173 | 1,846 | 1,862 | 155.8 | 46.3 | 22.2 | 670.7 | 76.0 |
| Kula | 471 | 560 | 520 | 118.9 | 47.4 | 20.8 | 444.9 | 82.8 |
| Lahaina | 70 | 388 | 369 | 136.5 | 43.3 | 23.4 | 840.1 | 72.2 |
| Pepeekeo | 37 | 3,376 | 1,322 | 144.8 | 58.8 | 22.6 | 812.3 | 71.3 |
| Poamoho | 209 | 911 | 770 | 130.1 | 52.7 | 22.3 | 656.2 | 76.7 |
| Wailua | 149 | 2,019 | 857 | 126.6 | 64.0 | 22.2 | 702.1 | 74.8 |

ET = Evapotranspiration; Net Rad = Net Radiation; Cloud Freq = Cloud Frequency; Temp = Average Annual Air Temperature; VPD = Vapor Pressure Deficit; RH = Relative Humidity

The trial was established in 2016 and the sites were selected to represent environmental parameters that could affect breadfruit production, including temperature, sunlight, wind, rainfall, salt spray, and soil type. All sites were managed in similar ways, with inter-row grasses mowed regularly, fertilization every 6 months (100 lbs 10-5-20 plus micronutrients, and 25 lbs of dolomite), and annual pruning in February once trees exceeded 12 feet in height.

### Gas-exchange measurements

At each site, leaf gas-exchange measurements were performed using a LI-COR 6800 portable photosynthesis system with the 6800-01A fluorometer light source (LI-COR Biosciences Inc., Lincoln, NE, USA). We selected mature leaves in positions two or three from the terminal end of a branch exposed to the sun (Lincoln et al., 2019b). For each tree, two leaves were measured in the intermediate lobes of the leaf, between lobes two and four from the leaf apex. For each variety, between three and six trees were measured at each site depending on differences in tree establishment related to site conditions. In particular, breadnut had difficulties in establishment and some sites do not include any data. For instantaneous measurements, conditions inside the LI-COR 6800 cuvette were controlled as follows: reference carbon dioxide concentration ([CO_2_]) at 400 ppm, photosynthetically active radiation (PAR) at 1000 mmol photons m^-2^ s^-1^ and flow rate at 600 mmol s^-1^. The vapor pressure deficit (VPD, kPa) and the leaf temperature (T_leaf_, °C) were allowed to vary with ambient conditions to expedite the stabilization of each measurement, which was recorded between two and three minutes after cuvette attachment. All measurements were made between 09:00 and 13:00 in March and April of 2018.

The light response curves were made with the same equipment and leaf selection as above, but the time of day, variety selection, and cuvette conditions were different. The PAR response curves were made in the afternoon, between 13:00 and 16:00 at all sites. The number of response curves per site varied depending on the sampling duration at each location: LAHA – 1 curve, KULA – 5, POAM – 2, WAIL – 2, PEPE – 2, KOHI – 1. All response curves were performed on the Ma’afala variety. Upon attachment of the cuvette, the leaf was allowed to stabilize under preset conditions for 30 minutes as follows: [CO_2_] at 400 ppm, PAR at 2000 mmol photons m^-2^s^-1^, flow rate at 600 mmol s^-1^, T_leaf_ between 30 and 35 ^°^ ^C^ (T_leaf_ was maintained constant during the response curve, but the exact value was based on ambient conditions), and VPD between 1.0 and 2.0 kPa (VPD was kept constant during the response curve, but exact VPD was based on ambient conditions). Once leaf stabilization occurred, an automated program was initiated to record measurements within three minutes of each step change in PAR. The PAR conditions were changed as follows: 2000, 1500, 1000, 800, 600, 500, 400, 300, 200, 100, 50, 0.

Measurements of the CO_2_ response curves were made as above, but site location and CO_2_ parameters were different. Given time constraints at each site, we chose to evaluate the CO_2_ response data only at the Waimanalo research site, which is not included in the ongoing variety trial. Given the earlier establishment of the site, the canopy size of Ma’afala trees were significantly larger than at the variety trial locations. However, branch positions exposed to the sun and leaf selection remained consistent. An automated program was used to record measurements at the following settings [CO2]: 400, 300, 200, 100, 50, 0, 400, 500, 600, 700, 800, 900, 1000, 1200, 1400, 1600, 2000.

The diurnal response curves were also made at the Waimanalo research site. These measurements were made in the same manner as the instantaneous measurements above, but leaf selection and frequency were different. ‘Sun leaves’ were selected based on their sun-exposed position on the southern facing aspect of mature trees in the plantation, where the external PAR sensor recorded maximum and minimum values of 2175 (midday) and 5 (evening) μmol photons m^-2^ s^-1^. ‘Shade leaves’ were also selected in the southern facing aspect, but were embedded within the tree canopy and the external PAR sensor recorded maximum and minimum values of 226 (midday) and 3 (evening) μmol photons m^-2^ s^-1^. We did record instances where a sun fleck penetrated the canopy and the values of PAR in shade leaves exceeded 1000 μmol photons m^-2^ s^-1^, but these were not considered normal conditions. We continually measured leaf function before direct sunlight and afterward, from 06:40 to 18:50 on a single day. We measured three Ma’afala trees in continuous progression throughout the day, so that the sun and shade leaves of the same tree were measured in direct succession, and each tree was measured once per hour. We have combined the data across trees, so all leaves measured within a 30-minute moving window are averaged together to provide a mean value.

### Additional plant measurements

Additional plant parameters measured in concert with gas-exchange included leaf stomatal density (mm^-2^), leaf mass by area (g cm^-2^), leaf nitrogen (N) content (%), leaf SPAD, and leaf fluorescence. Tree diameter growth rates (cm y^-1^) and breadfruit yields (kg y^-1^) were obtained as part of ongoing trial monitoring.

Stomatal density was measured using clear nail polish imprints on the abaxial surface of the leaf. Nail polish was applied to the central portions on the third or fourth lobe from the leaf tip, away from the central vein and leaf edge; each leaf was in the second or third position from the apex of a sun-exposed mid-canopy branch. Two leaves per tree were collected. Once the nail polish was dried, it was removed using clear adhesive tape and attached to glass microscope slides. The microscope slides were imaged using a Nikon Eclipse NiE brightfield microscope (Nikon Corporation, Tokyo, Japan) with a mounted DS-Qi1 monochrome cooled digital camera and the 10x objective (image field of view was 819 by 655 um). Stomatal density was manually counted on four images per imprint using the cell counter in Fiji (Schindelin et al. 2012). The average stomatal density was calculated across all images and leaves per tree. Due to sampling constraints, stomatal density was only quantified at three sites. WAIL, KOHI, and PEPE. Although breadnut imprints and images were collected from the three sites, they were only quantifiable from PEPE because of the quality of the image.

The mass area and N-content were measured by taking 25 25mm diameter leaf punches from three leaves of each tree. The punches were distributed throughout the leaf and avoided large veins. The leaf punches were weighed wet, dried at 60 °C, and again weighed dry to determine the mass of a known area of leaf material. After weight determination, the punches were ground and the nutrient content was analyzed at Brookside Laboratory (New Bremem, Ohio) according to the same procedures as Lincoln *et al. (*2019a).

SPAD and leaf fluorescence were measured with a 502 SPAD plus meter (Konika Minolta, Tokyo, Japan) and a PPM 300 plant photosynthetic meter (Environmental Analysis and Remote Sensing Earth Environmental Monitoring, Delft, The Netherlands). Ten leaves in the second or third position from the apex of sun-exposed mid-canopy branches were measured and averaged from each tree.

The growth rates and fruit yields were measured by site-specific technicians. The diameter at breast height was measured twice a year since the start of the trial, taking the diameter 1.3 m from the ground surface using a cloth diameter tape. Yields were monitored every two weeks. All male flowers and fruit of all stages were counted for each tree. All mature fruit were harvested, counted, and weighed for each tree. The total yield was calculated as the sum of the yield per tree for the calendar year, averaged among individuals of the same variety at each location.

### Statistical analysis

Carbon assimilation data from PAR response curves were fitted to a nonlinear model as described by Lobo et al. (2013), which can be used to derive the quantum yield, light-saturated photosynthesis, light saturation point, light compensation point, dark respiration rates and 90% light saturated photosynthesis for each curve. Carbon assimilation data from CO_2_ response curves were fitted to the FvCB model of C3 photosynthesis (Farquhar et al. 1980) using the tools provided in Sharkey (2016). Water-use efficiency was calculated as the ratio of net carbon assimilation to stomatal conductance under ambient conditions. ANOVA and Tukey’s HSD tests of comparative means were used to assess statistical differences between groups (i.e., sites, varieties, site x variety). Linear and nonlinear regressions were performed to examine the relationships between variables. All statistical analyses were performed in JMP (SAS Institute; Cary, NC), RStudio (PBC, Boston, MA), or Microsoft Excel.

## Results

### Carbon assimilation by site and variety

Net photosynthesis exhibited significant variability according to site (p < 0.001) and variety (p < 0.001), but there was no interactive effect between site and variety (p = 0.429). When assessing site-specific differences, separation in assimilation rates was evident, ranging from a mean (and standard error) of 12.28 (±0.59) to 20.25 (±1.05) µmol m⁻² s⁻¹ (Figure 2a). The variation demonstrated a clear distinction between highly productive sites, characterized by robust carbon assimilation, and low productivity sites. This photosynthetic distinction is underscored by wide differences in fruit yield across sites, as illustrated in Figure 1. These initial observations led us to categorize the sites into ‘high’ and ‘low’ productivity to facilitate the presentation, interpretation, and discussion of the data. Throughout the results, we maintain a consistent coloration of red as representing the high-productivity sites and blue as representing the low-productivity sites.

**Figure 2:**
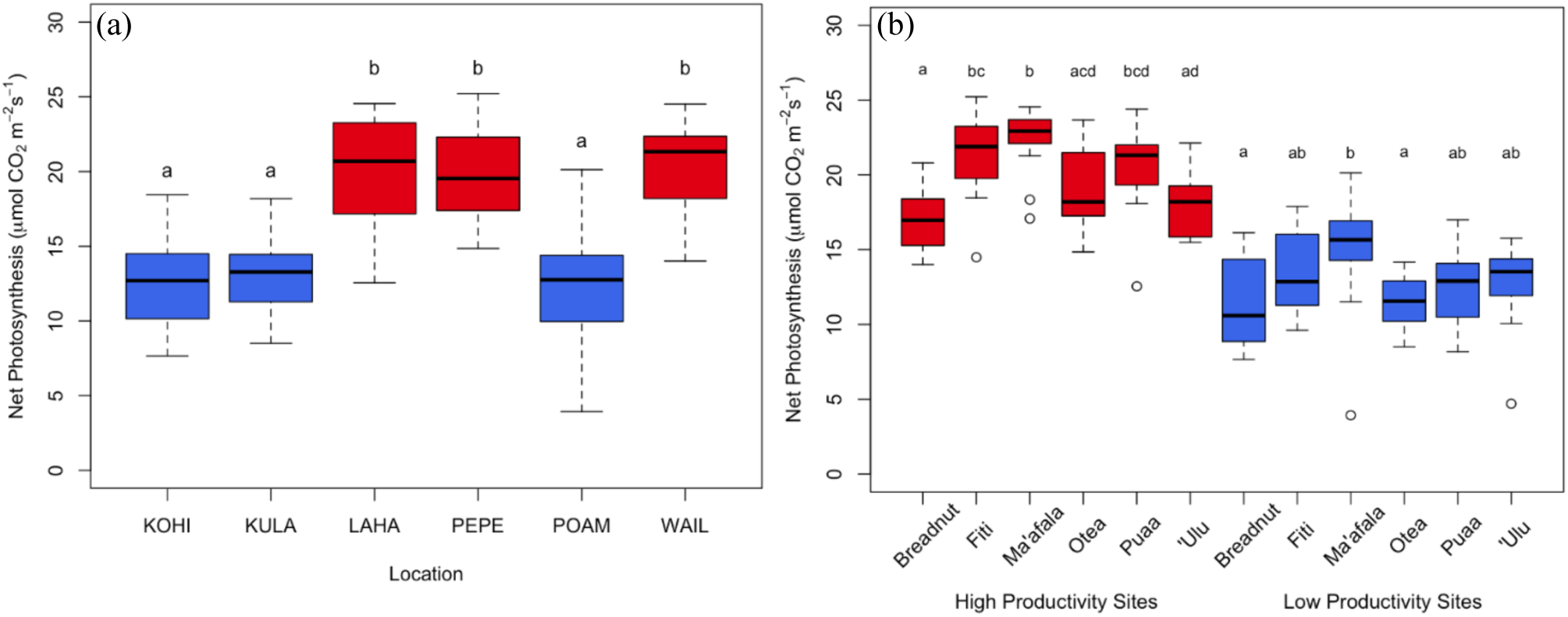
Carbon assimilation rates derived from instantaneous gas-exchange measurements categorized by (a) site, and (b) site and variety. Red indicates measurements at high-productivity sites and blue indicates measurements at low-productivity sites; letters indicate significant differences between categorial groups (p<0.05).

Variety analysis demonstrated subtle differences in the rate of assimilation between varieties, with some statistical differences (Figure 2b). These separations were more pronounced at high-productivity sites, suggesting that the superior varieties have the potential to thrive in environments conducive to increased photosynthesis. At all sites, Ma’afala consistently exhibited the highest assimilation rates among all varieties, while breadnut consistently displayed the lowest rates, although these were not significantly different from the next closest variety.

### Response to light intensity and carbon dioxide

Analysis of the light response curves indicated variations based on site classification, again distinguishing between high-and low-productivity sites (Figure 3). The derivation of critical parameters was based on the average values at each group of sites, represented by dark-filled squares, fitted to a nonlinear model (Lobo et al., 2013). These parameters provide an opportunity for a direct comparison of light-dependent photosynthetic traits at high-and low-productivity sites (Table 2). For example, the light saturation point was notably higher with high-productivity sites at a mean of 1545 µmol m⁻² s⁻¹ compared to a mean of 1183 µmol m⁻² s⁻¹ at low-productivity sites. Meanwhile, carbon respiration and the light compensation point were higher at low-productivity sites.

**Figure 3:**
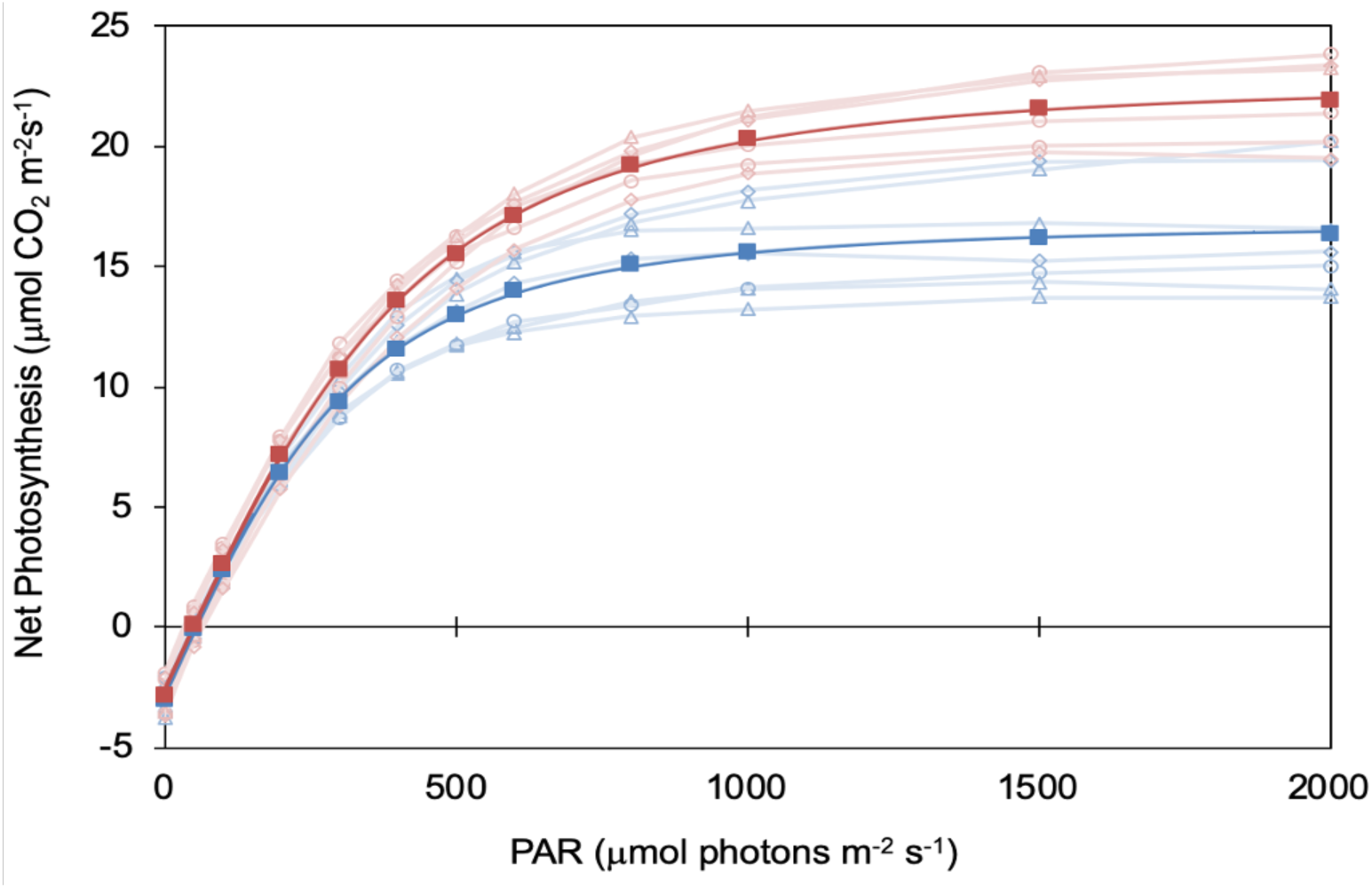
Carbon assimilation response curves to photosynthetically active radiation (PAR), separated by high-(red) and low-productivity sites (blue). The mean response curve across high-and low-sites, shown with bold color and filled squares, was fitted using the photosynthetic model parameterization of Lobo *et al*. (2013). The individual unfitted response curves are shown in light colors and unfilled objects (red circle = WAIL, red triangle = LAHA, red diamond = PEPE; blue circle = KOHA, blue triangle = KULA, blue diamond = POAM). All response curves were performed only on Ma’afala.

**Table 2:** Photosynthetic parameters derived from the average light response curve across low-(LPS) and high-productivity sites (HPS), using the model parameterization from Lobo *et al*. (2013).

| Parameter | LPS | HPS | Units |
| --- | --- | --- | --- |
| $P_{\text{gmax}}$ | 19.7 | 25.4 | $\mu\text{mol CO}_2 \text{ m}^{-2}\text{s}^{-1}$ |
| $P_{\text{N}(I_{\text{max}})}$ | 15.9 | 21.6 | $\mu\text{mol CO}_2 \text{ m}^{-2}\text{s}^{-1}$ |
| $R_{\text{D}}$ | 2.9 | 2.7 | $\mu\text{mol CO}_2 \text{ m}^{-2}\text{s}^{-1}$ |
| $\phi_{(I_0)}$ | 0.0533 | 0.0528 | $\mu\text{mol CO}_2 \mu\text{mol}^{-1} \text{ photons}$ |
| $I_{\text{comp}}$ | 54.2 | 50.6 | $\mu\text{mol photons m}^{-2}\text{s}^{-1}$ |
| $I_{\text{sat}(50)}$ | 258 | 319 | $\mu\text{mol photons m}^{-2}\text{s}^{-1}$ |
| $I_{\text{sat}(90)}$ | 835 | 1060 | $\mu\text{mol photons m}^{-2}\text{s}^{-1}$ |
| $I_{\text{max}}$ | 1183 | 1545 | $\mu\text{mol photons m}^{-2}\text{s}^{-1}$ |

$P_{\text{gmax}}$ = maximum gross photosynthesis rate; $P_{\text{N}(I_{\text{max}})}$ = maximum net photosynthesis rate at $I_{\text{max}}$ ; $R_{\text{D}}$ = dark respiration rate; $\phi_{(I_0)}$ = quantum yield at $I = 0$ ; $I$ = photosynthetically active radiation in photons; $I_{\text{(comp)}}$ = light compensation point; $I_{\text{sat}} (50)$ = light saturation point at 50% of $P_{\text{N}(I_{\text{max}})}$ ; $I_{\text{sat}(90)}$ = light saturation point at 90% of $P_{\text{N}(I_{\text{max}})}$ ; $I_{\text{max}}$ = light saturation point of $P_{\text{N}(I_{\text{max}})}$

Response curves to increasing carbon dioxide concentration (traditional *A*-*C*_i_ curves) were only performed on the Ma‘afala variety at the Waimanalo site and were fitted to a biochemical model of C3 photosynthesis (Farquhar et al. 1980, Figure S1). These curves allowed for an initial estimate of the biochemical limitations of photosynthesis in breadfruit (reported as the mean and standard error of four replications). The maximum rate of carboxylation of the Rubisco enzyme (*V*_cmax_) was estimated at 151 (±20.8) μmol m^-2^s^-1^ and limited photosynthesis at approximately 400 ppm ambient CO_2_ and below; the maximum rate of electron transport (*J*_max_) was 128 (±12.2) μmol m^-2^s^-1^ and limited photosynthesis between 400 and 1000 ppm CO_2_; and the maximum rate of use of triose phosphate (*TPU*) was 8.3 (±0.8) μmol m^-2^s^-1^ and limited photosynthesis above 1000 ppm CO_2_. Consequently, the saturation point for breadfruit photosynthesis under increasing ambient CO_2_ and otherwise nonlimiting conditions should be approximately 1000 ppm.

### Diurnal patterns and shade leaves

The diurnal response curves revealed substantial oscillations in carbon assimilation influenced by stomatal function (Figure 4) in mature Ma’afala trees at the Waimanalo site. Leaves exposed to full sunlight showed oscillations in the assimilation rate from full maximum to completely zero early in the day, with moderation of the oscillations occurring in the afternoon as stomatal function stabilized. These patterns were driven by diffusion limitations in the stomatal conductance of sun-exposed leaves, while shade leaves exhibited only mild oscillations in comparison. Noticeably, photosynthesis and stomatal conductance reached maximum values during the first peak, but stomatal conductance showed a lower percentage of recovery (49%) compared to photosynthesis (74%), suggesting a mechanism to limit water loss while maintaining adequate photosynthesis. This differential recovery improved the intrinsic water-use efficiency from 38.3 μmol CO_2_ mol^-1^ H_2_O around 9 am to 57.3 μmol CO_2_ mol^-1^ H_2_O around 11:30 am, representing a 50% increase. Shade leaves had significantly lower photosynthetic rates than sun leaves throughtout the day, with mean (and standard error) of 5.77 (±0.56) and 9.08 (±1.08) µmol CO_2_ m⁻² s⁻¹ for shade leaves (n = 36), and sun leaves (n = 36) respectively, even under the same instantaneous PAR conditions in the gas-exchange chamber (1000 µmol photons m⁻² s⁻¹).

**Figure 4:**
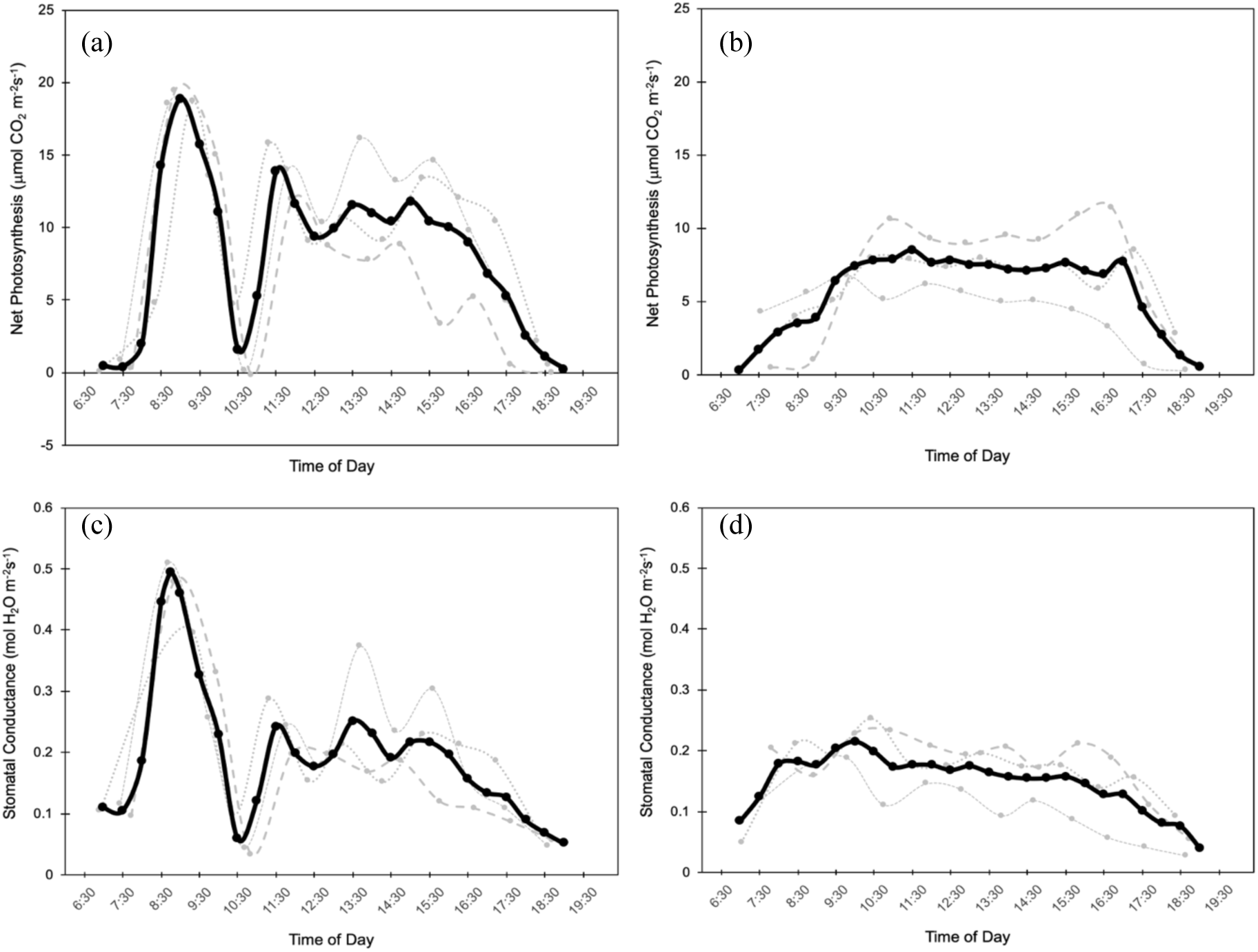
Diurnal response curves of carbon assimilation (a, b) and stomatal conductance (c, d) for sun leaves (a, c) and shade leaves (b, d) over the course of sunlight hours in a single day. The response curves of individual leaves are shown in gray (one per tree, distinguished by filled circles and smoothed connecting dots, short dashes, or long dashes), while the 30-minute moving average of all leaves is shown in black (filled circles and smoothed line).

### Stomatal density and water-use efficiency

Stomatal density exhibited variation between varieties (Figure 5a) and showed a significant correlation with net photosynthesis (Figure 5b). Although there were statistical differences between varieties (p < 0.001), variation in stomatal density within a variety was also substantial, ranging between 200 – 400 stomata per mm^2^ in maximum and minimum replicates. There was no location-specific effect on stomatal density (p = 0.100), but there was an interactive effect between location and variety (p < 0.05). The relationship between stomatal density and carbon assimilation was positively correlated for both high-and low-productivity sites and when averaged across all sites. There is a very tight correlation and a steeper slope at the low-productivity site (KOHI Figure 5b, r^2^ = 0.98, p < 0.001, slope = 0.042 μmol CO_2_ m^-2^s^-1^ per stoma), indicating that constraints on leaf diffusion could limit photosynthesis at this location. In contrast, a weaker relationship at high-productivity sites suggests that factors other than stomatal diffusion might also limit photosynthesis in these environments (WAIL and PEPE Figure 5b, r^2^ = 0.37, p < 0.05, slope = 0.018 μmol CO_2_ m^-2^s^-1^ per stoma). When comparing all high-and low-productivity sites, intrinsic water-use efficiency was not statistically distinguishable, with means (and standard error) of 61.2 (±2.2) and 57.2 (±2.2) μmol CO_2_ mol^-1^ H_2_O, respectively.

**Figure 5:**
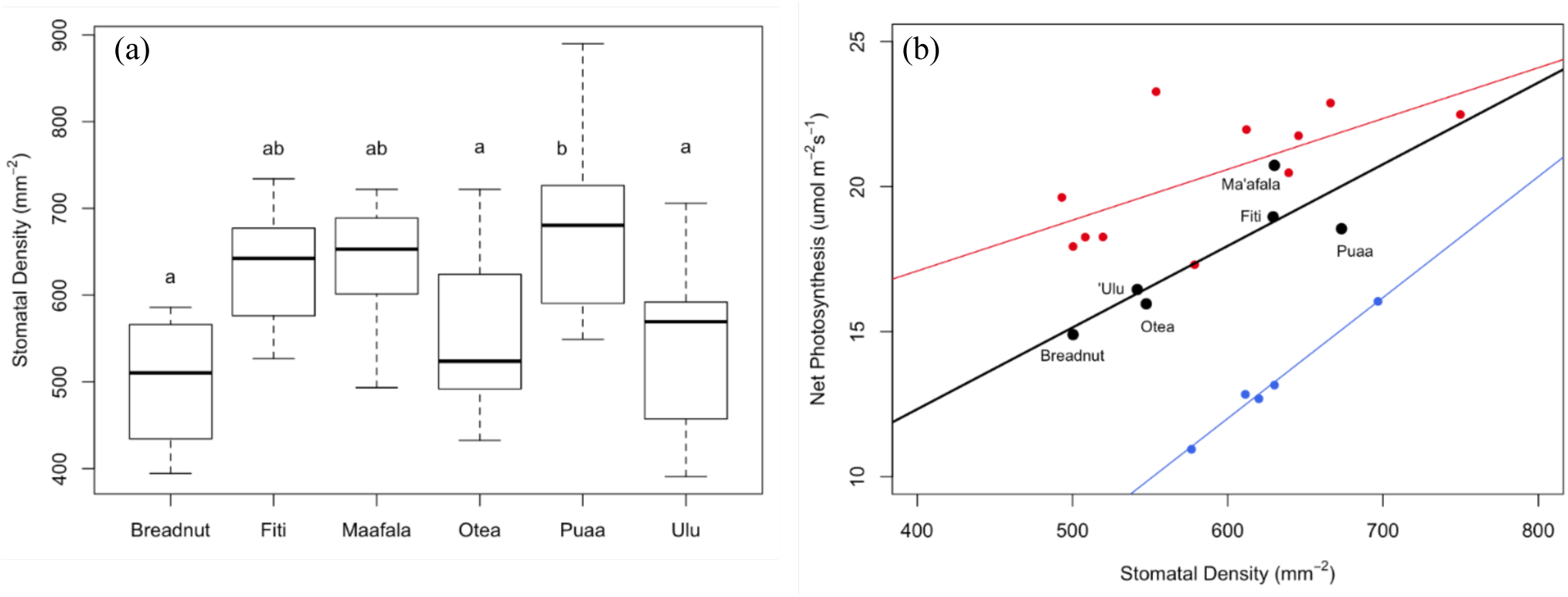
Stomatal density of different varieties at three sites (a) and correlated with carbon assimilation at high-and low-productivity sites (b). Letters above boxplots in (a) indicate significant differences between varieties when combining data from sites WAIL, PEPE, and KOHI. In (b), red circles and regression line indicate all varieties at high-productivity sites WAIL and PEPE, blue circles and regression line indicate all varieties at low-productivity site KOHI, and black circles and regression line indicate all varieties at WAIL, PEPE, and KOHI combined. Each circle represents the average value of the variety.

### Photosynthesis, leaf nitrogen content, tree growth, and fruit yields

Leaf mass area (g cm^-1^), leaf nitrogen content (%), SPAD, and fluorescence measurements generally differed between sites but not between varieties (Table S4). At all sites, photosynthesis had a significant but weak relationship with leaf N measured per unit mass (r^2^ = 0.38, p < 0.001) or per unit area (r^2^ = 0.22, p<0.005). However, when examining the relationship within low and high-productivity sites, neither showed a statistical relationship with leaf N (Figure 6), suggesting that in both cases, nitrogen is not the limiting element in photosynthesis. Carbon assimilation per unit N differed significantly between the two groups, with means (and standard error) of 587 (±34) and 724 (±33) µmol CO_2_ gN^-1^ s⁻¹ for low and high-productivity sites, respectively. The percentage of leaf N was strongly correlated with the SPAD measurements, across all samples (r^2^ = 0.82, p < 0.001) and when separated by low-(r^2^ = 0.77, p < 0.001) and high-(r^2^ = 0.65, p < 0.001) productivity sites.

**Figure 6:**
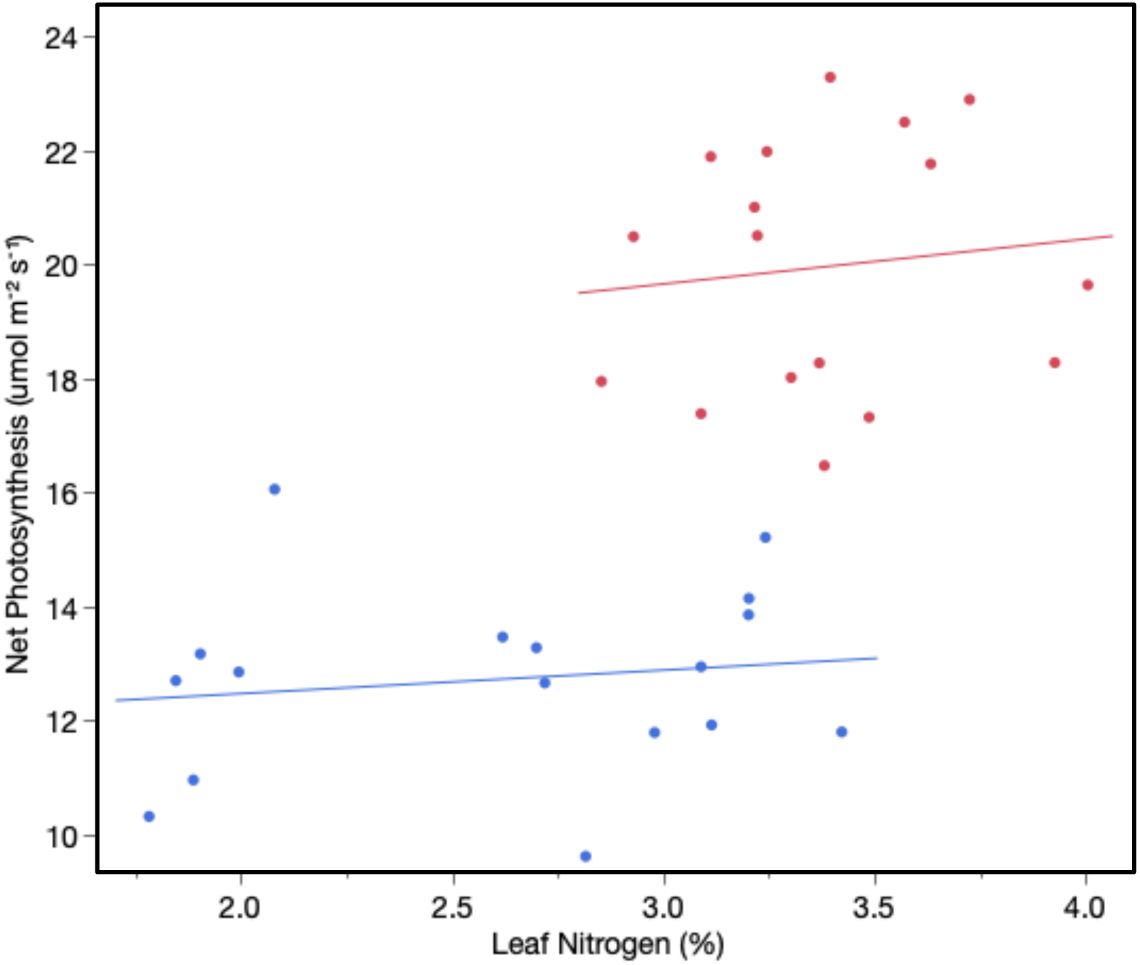
Relationship between leaf N concentration and carbon assimilation at high-and low-productivity sites. Each point represents the average value for each variety at an individual site (n=17 per group; breadnut did not establish adequately at two sites, one per group), with red circles for high-productivity sites and blue circles for low-productivity sites.

Assimilation rates show a strong positive correlation between tree growth and productivity at all sites measured by diameter at breast height (r^2^ = 0.84, p < 0.001) and total fruit weight produced in a year (r^2^ = 0.80, p < 0.001). When separated by sites of high-and low-productivity, significant positive relationships of growth (r^2^ = 0.20, p < 0.05; r^2^= 0.28, p < 0.05) and yield (r^2^ = 0.46, p < 0.005; r^2^ = 0.24, p < 0.05) were still observed, although the relationships were weaker (Figure 7). The regression slope indicates that for each additional unit of carbon assimilation (1.0 µmol CO_2_ m⁻² s⁻¹) an additional 17.8 kg of fruit is produced per year at high-productivity sites compared to 6.1 kg of fruit at low-productivity sites.

**Figure 7:**
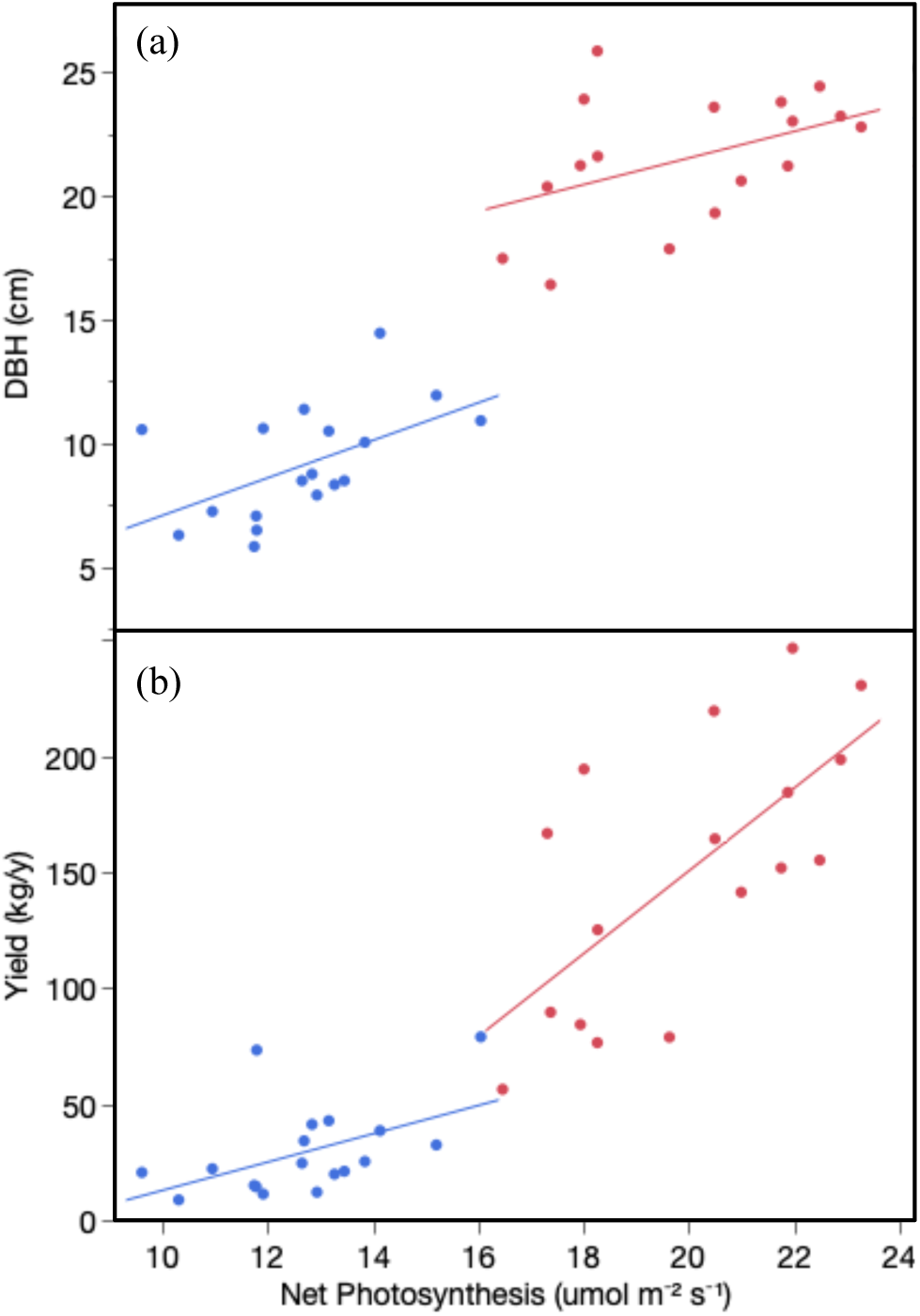
Relationship between carbon assimilation and tree size (a) and fruit yield (b). Each point represents the average variety value at an individual site (n=17 per group; breadnut did not establish adequately at two sites, one per group), separated by high-(red) and low-productivity sites (blue).

## Discussion

This study explored the photosynthetic parameters associated with different varieties of breadfruit grown in multiple environments. To our knowledge, this is the first comprehensive investigation of breadfruit carbon assimilation, associated leaf function, and impacts on growth and yields. Here, we discuss these findings in terms of the potential adaptation of breadfruit from a predominantly forest-dwelling species to one that can be used in various agricultural systems.

Breadfruit demonstrated a light saturation point that was comparable to other temperate and tropical fruit tree crops, achieving light saturation at 1183 and 1545 µmol m⁻² s⁻¹ in low-and high-productivity sites, respectively (Table 2). In other tree fruit species, these values tend to vary across a wide spectrum, with some subtropical species below breadfruit values, including 750 – 1000 in orange (*Citrus sinensis*) and 1100 in fig (*Ficus carica*), while papaya (*Carica papaya*) required 1900 µmol m⁻² s⁻¹ for complete light saturation (Restrepo-Diaz et al. 2010). This variation is also true in temperate fruit tree species that are almost exclusively grown in full-sunlight orchard systems, ranging from 700 – 800 in pecan (*Carya illinoinensis*), to 1300 in peach (*Prunus persica*), and finally 1800 – 1900 µmol m⁻² s⁻¹ in apple trees (*Malus domestica*) (Restrepo-Diaz et al. 2010).

While breadfruit has traditionally been cultivated in forest-based agricultural systems with lower amounts of direct sunlight, it appears to possess leaf photochemistry that is very capable of harnessing extra light to enhance carbon assimilation. Furthermore, this biochemical capacity of breadfruit was corroborated by enzymatic responses to increasing ambient CO_2_ (Figure S1), where estimated values of *V*_cmax_ (151.4 μmol m^-2^s^-1^) and *J*_max_ (127.5 μmol m^-2^s^-1^) are at the higher end of previously reported data, both within tropical forest species (*V*_cmax_ ranging between 9 – 126 and *J*_max_ ranging between 30 – 222) and horticultural fruit trees (*V*_cmax_ ranging between 11 – 150 and *J*_max_ ranging between 29 – 148, including peach (*Prunus persica*), grapefruit (*Citrus paradisi*), and lemon (*Citrus limon*)) (Lloyd et al. 1992; Wullschleger 1993).

These initial results are promising with respect to the transition to orchard-based systems, with two caveats being varietal choice and site suitability. All photosynthetic response curves were performed only on the Ma’afala variety, which had the highest rates of carbon assimilation between varieties, regardless of location (Figure 2). Thus, the photochemical potential in other varieties may be further reduced and should be investigated to determine if a varietal other than Ma’afala is preferred. However, varieties like Fiti and Puaa had photosynthetic values that were comparable to Ma’afala and should provide similar yield returns (Figure 2 and Table S2). In terms of general physiological comparisons between varieties, it was unexpected that the varieties ‘Ulu and Otea were closer to Breadnut than to Ma’afala and Fiti. ‘Ulu and Otea are eastern Pacific varieties that have presumably undergone more selection as the Polynesian people moved from west to east. Across all varieties, there was still a high degree of parameter variability, including net photosynthesis and stomatal density, even within a variety. This may be indicative of a less intense selection of functional traits over time, as targeted backcrossing and inbreeding tend to reduce variability. The discussion of breeding and selection is also quite complicated in the case of breadfruit, as the varieties were often clonally propagated through ‘root runners’ to ensure phenotypic consistency and current varieties can be fertile diploids or infertile triploids.

The second caveat, in terms of site suitability for orchard-based systems, was also obvious when comparing photosynthetic responses to fruit yields and growth rates (Figures 1 and 2, Table S2 and S3). There were clear differences between low-and high-productivity sites that informed our data analysis, where carbon assimilation differed from 12.3 to 20.3 µmol CO_2_ m⁻²s⁻¹ and fruit yield differed from < 50 to between 100 and 200 kg/y, respectively (Figure 7). Although these performance differences are evident, unfortunately there has not been a singular environmental parameter or site characteristic that can explain it (Lincoln et al. 2019a). As such, we believe the differences are likely due to the combination of multiple variables acting at each site, which makes the isolation of any one parameter difficult. Our previous assessment of five variety trial locations suggested that elevation and solar radiation were the most influential environmental parameters on the early growth of breadfruit (Lincoln et al. 2019a). However, sunlight is not a limiting factor in tropical ecosystems, and leaves of evergreen tropical trees can generally achieve these photosynthetic rates at lower photon flux densities (Jayasekara and Jayasekara 1995). This is certainly true for breadfruit in our study, whereby 90% photosynthetic saturation occurred at or below 1060 PAR, which is less than half the maximum midday PAR observed, 2175 µmol photons m⁻² s⁻¹. As there was almost complete overlap at the sites between the studies, the negative relationship with elevation remained consistent in this study. The three low-productivity sites had higher elevations than the high-productivity sites, with average elevations of 285 and 85m, respectively (Table 1). How elevation might determine productivity remains to be verified. In the context of leaf physiology, it could be considered that elevation increases exposure to UV radiation and, in combination with excessive tropical sunlight, physically damages the molecular photosystems or plant cells of the leaf (Shi and Liu, 2021). This potential hazard deserves further scientific investigation and consideration in possible plantation schemes and site location choices.

Although the breadfruit photosynthetic apparatus appears capable of using full sunlight in plantation schemes at preferred locations, we also observed a very dramatic pattern of stomatal oscillation in mature trees at Waimanalo (Figure 4). Our initial results from a complete set of diurnal measurements of carbon assimilation and stomatal conductance indicate that this is likely a water savings or hydraulic protection mechanism, which increases intrinsic water-use efficiency by 50%. Observations of stomatal oscillation have been found in other subtropical and tropical fruit trees, including orange (*Citrus sinensis*) and banana (*Musa acuminata*), so this phenomenon is not distinct from breadfruit (Dzikiti et al. 2007; Zait et al. 2017). However, a similarly strong oscillation pattern did not occur in the shade leaves of the same tree nor in the sun leaves of the younger breadfruit trees observed throughout the variety trial. Waimanalo was planted two years before multi-environmental sites and the Ma’afala trees had a much larger average canopy volume than even the fastest-growing variety trial location in Wailua (18.8 m^3^ to 3.72 m^3^ in volume at the time of measurement, respectively). Given that limitations on canopy size are removed in plantation schemes and the vast surface area of meter-long breadfruit leaves, it appears that the native water delivery network – either the root systems, trunk and branch conduits, or leaf vasculature – is not sufficient to deliver the required amount of water transpired in full sunlight across such a broad canopy (Table S1; Buckley, 2005). Consequently, a potentially very negative water potential can develop in sun leaves that invokes a mechanical or osmotic stomatal response to limit water loss. The threat of hydraulic failure and the need for sufficient water resources must also be considered in breadfruit plantation schemes. Novel approaches to thinning the leaf canopy must also be explored to limit tree size, increase planting density, or reduce per tree water requirements.

Finally, breadfruit assimilation showed a tight coupling with stomatal density (Figure 5b) and stomatal conductance (Figure 4). The correlation of the former relationship also differed in strength between the high-and low-productivity sites. This may indicate that breadfruit assimilation is predominantly limited by CO_2_ supply rather than photosynthetic biochemistry (Figure 3 and Figure S1). That is, the diffusion of CO_2_ from outside the leaf to carboxylation sites in the chloroplasts is more limiting than the Rubisco capture of CO_2_ and subsequent photochemistry. Given the hypostomatous nature and large size of the breadfruit leaves, limitations to diffusion are not unexpected. An initial estimate of internal mesophyll conductance (*g*_m_) was equal to 0.9 (± 0.1) μmol CO_2_ m^-2^s^-1^pa^-1^ (Figure S1). That value places breadfruit at the lower end of the published *g*_m_ in subtropical and tropical tree fruits (4.1 ± 0.2 in peach (*Prunus persica*), 2.2 ± 0.3 in grapefruit (*Citrus paradisi*), 1.7 ± 0.1 in lemon (*Citrus paradisi),* and 1.1 ± 0.1 in macadamia (*Macadamia integrifolia*); Lloyd et al., 1992). This internal limitation of CO_2_ transfer can potentially be overcome by increasing diffusion pathways through stomatal density, especially at low-productivity sites where the opening of stomata may be limited by environmental conditions (Figure 5b). However, the negative feedback of increasing stomatal conductance on plant water status, stomatal oscillations, and the related temporal limitations on carbon assimilation remain a concern at high-productivity sites under full sunlight (Figure 4). Given that biochemical efficiency and nitrogen content do not appear to be limiting factors for carbon assimilation (Figures 3 and 6, Figure S1), water transport and CO_2_ diffusion may be the dominant physiological factors that restrict productivity in potential breadfruit plantation schemes.

## Supplementary data

*Figure S1:* Photosynthetic response curves to changes in ambient carbon dioxide concentration.

*Table S1:* Stomatal conductance (mol H_2_O m^-2^s^-1^) by site and variety.

*Table S2:* Fruit yield (kg per year) by site and variety.

*Table S3:* Tree trunk diameter at breast height (cm) by site and variety.

*Table S4:* Average values of leaf mass by area, leaf nitrogen content, SPAD, and fluorescence by site and variety.

## Acknowledgements

The authors would like to thank each of the host sites and associated staff for trial upkeep and help with data collection, and various members of the Indigenous Cropping Systems Laboratory for help with data collection and sample analysis.

## Author Contributions

NL established the variety trial; NL and GD designed the experimental and measurement approaches; NL, GD and DA collected and analyzed the data; NL and GD wrote the manuscript; NL, GD, DA, and RP edited the manuscript.

