## Supplementary Data for "From forests to farming: identification of photosynthetic limitations in breadfruit across diverse environments"

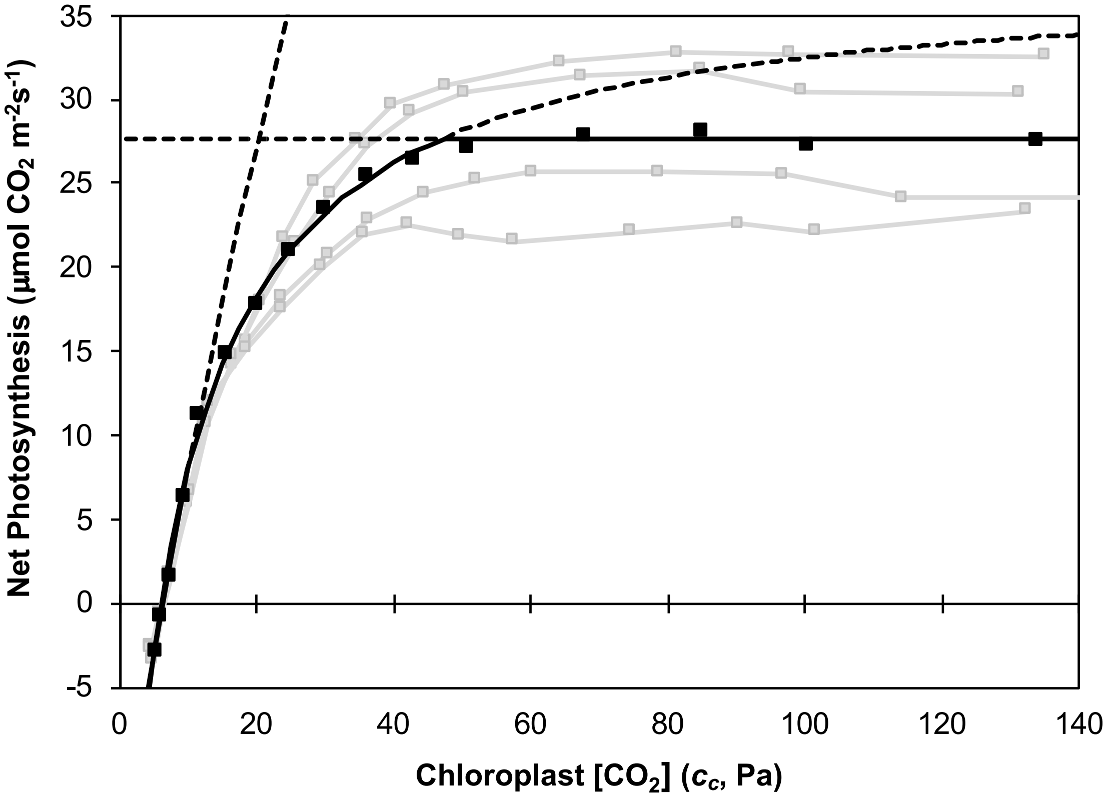


**Figure S1: Photosynthetic response curves to changes in ambient carbon dioxide concentration.** Gray squares and lines are four individual response curves of the Ma'afala variety, while black squares and soild lines are a representative curve of average values fitted with the FvCB biochemical model of C3 photosynthesis. The solid black line is the combination of three separate curves that define different stages of biochemical limitation – initially Rubisco enzyme kinetics, then RuBP regeneration, and finally, end product formation – while the dashed black lines are the extension of those curves in $CO_2$ regions where they are not limiting. The following photosynthetic parameters were defined using the model fit for each gray replica curve, presented here as the mean (and standard error): $V_{cmax}$ = 151.4 (20.8) μmol $m^{-2}s^{-1}$; $J_{max}$ = 127.5 (12.2) μmol $m^{-2}s^{-1}$; *RDU* = 8.3 (0.8) μmol $m^{-2}s^{-1}$; $R_d$ = 3.3 (0.4) μmol $m^{-2}s^{-1}$; $g_m$ = 0.9 (0.1) μmol $m^{-2}s^{-1}pa^{-1}$; where $V_{cmax}$ is the maximum carboxylation rate of Rubisco, $J_{max}$ is the maximum electron transport for the given light intensity (1000 μmol photons $m^{-2}s^{-1}$), *TPU* is the maximum rate of use of triose phosphate, $R_d$ is daytime respiration, and $g_m$ is the mesophyll conductance to internal $CO_2$ transfer.

Table S1: Stomatal conductance (mol $H_2O$ $m^{-2}s^{-1}$) by site and variety.

| | WAIL | PEPE | LAHA | KOHI | KULA | POAM | Mean | SE |
|---|---|---|---|---|---|---|---|---|
| **Ma'afala** | 0.463 | 0.468 | 0.448 | 0.343 | 0.212 | 0.222 | **0.359** | 0.045 |
| **Fiti** | 0.404 | 0.392 | 0.348 | 0.265 | 0.177 | 0.214 | **0.300** | 0.036 |
| **Otea** | 0.368 | 0.378 | 0.333 | 0.235 | 0.222 | 0.212 | **0.291** | 0.029 |
| **Puaa** | 0.415 | 0.313 | 0.270 | 0.288 | 0.187 | 0.170 | **0.274** | 0.033 |
| **'Ulu** | 0.313 | 0.328 | 0.294 | 0.228 | 0.242 | 0.147 | **0.259** | 0.025 |
| **Breadnut** | 0.309 | 0.300 | -- | 0.242 | 0.194 | 0.189 | **0.247** | 0.023 |
| Mean | **0.379** | **0.363** | **0.339** | **0.267** | **0.206** | **0.193** | | |
| SE | 0.025 | 0.026 | 0.031 | 0.018 | 0.010 | 0.012 | | |

Table S2: Fruit yield (kg per year) by site and variety.

| | WAIL | PEPE | LAHA | KOHI | KULA | POAM | Mean | SE |
|---|---|---|---|---|---|---|---|---|
| **Ma'afala** | 198.3 | 230.3 | 184.2 | 78.9 | 32.4 | 21.0 | **124.18** | 37.19 |
| **Fiti** | 151.7 | 246.3 | 141.2 | 42.8 | 25.2 | 19.8 | **104.49** | 36.94 |
| **Otea** | 78.8 | 166.6 | 164.2 | 22.1 | 73.2 | 14.3 | **86.53** | 27.10 |
| **Puaa** | 155.0 | 219.3 | 89.5 | 34.1 | 11.2 | 24.5 | **88.93** | 33.93 |
| **'Ulu** | 125.0 | 76.4 | 194.2 | 41.2 | 38.5 | 20.5 | **82.63** | 26.96 |
| **Breadnut** | 56.3 | 84.2 | 10.9 | 8.7 | 14.9 | 12.0 | **31.17** | 12.90 |
| Mean | **127.51** | **170.50** | **130.70** | **37.97** | **32.58** | **18.69** | | |
| SE | 21.45 | 30.56 | 28.39 | 9.72 | 9.14 | 1.89 | | |

Table S3: Tree trunk diameter at breast height (cm) by site and variety.

| | WAIL | PEPE | LAHA | KOHI | KULA | POAM | **Mean** | **SE** |
|---|---|---|---|---|---|---|---|---|
| **Ma'afala** | 23.2 | 22.8 | 21.2 | 10.9 | 11.9 | 8.5 | **16.42** | 2.73 |
| **Fiti** | 23.8 | 23.0 | 20.6 | 10.5 | 10.0 | 8.3 | **16.04** | 2.92 |
| **Otea** | 17.8 | 20.4 | 19.3 | 7.2 | 6.5 | 7.0 | **13.04** | 2.76 |
| **Puaa** | 24.4 | 23.6 | 16.4 | 11.4 | 10.6 | 8.5 | **15.80** | 2.80 |
| **'Ulu** | 21.6 | 25.9 | 23.9 | 8.7 | 14.4 | 10.5 | **17.51** | 2.96 |
| **Breadnut** | 17.5 | 21.2 | 5.8 | 6.3 | 5.8 | 7.9 | **10.74** | 2.78 |
| Mean | **21.39** | **22.80** | **17.86** | **9.16** | **9.88** | **8.46** | | |
| SE | 1.24 | 0.78 | 2.62 | 0.85 | 1.34 | 0.47 | | |

Table S4. Average values of leaf mass by area, leaf nitrogen content, SPAD, and fluorescence by site and variety.

| Site | Variety | Leaf Mass Area (g cm$^{-2}$) | Nitrogen (%) | SPAD | Fluorescence (%) |
|---|---|---|---|---|---|
| WAIL | Breadnut | 5.68 | 3.38 | 37.59 | 52.04 |
| WAIL | Fiti | 7.25 | 3.63 | 58.37 | 61.30 |
| WAIL | Ma'afala | 7.29 | 3.73 | 56.38 | 63.76 |
| WAIL | Otea | 7.29 | 4.01 | 60.18 | 51.56 |
| WAIL | Puaa | 7.43 | 3.57 | 53.14 | 64.76 |
| WAIL | 'Ulu | 7.41 | 3.93 | 57.24 | 50.20 |
| PEPE | Breadnut | 8.48 | 2.85 | 50.80 | 48.72 |
| PEPE | Fiti | 8.91 | 3.25 | 54.89 | 47.92 |
| PEPE | Ma'afala | 8.88 | 3.40 | 53.68 | 53.16 |
| PEPE | Otea | 9.06 | 3.49 | 55.56 | 48.88 |
| PEPE | Puaa | 9.53 | 2.93 | 49.84 | 50.00 |
| PEPE | 'Ulu | 9.35 | 3.37 | 54.36 | 46.88 |
| LAHA | Breadnut | 7.30 | 2.62 | 44.16 | 54.50 |
| LAHA | Fiti | 8.16 | 3.22 | 55.31 | 52.13 |
| LAHA | Ma'afala | 8.14 | 3.11 | 53.59 | 49.64 |
| LAHA | Otea | 8.61 | 3.22 | 58.80 | 46.75 |
| LAHA | Puaa | 8.52 | 3.09 | 52.81 | 38.93 |
| LAHA | 'Ulu | 8.42 | 3.30 | 51.31 | 49.50 |
| KOHI | Breadnut | 6.82 | 1.78 | 29.66 | 35.60 |
| KOHI | Fiti | 8.84 | 1.90 | 35.88 | 35.52 |
| KOHI | Ma'afala | 9.15 | 2.08 | 41.83 | 32.36 |
| KOHI | Otea | 8.99 | 1.89 | 30.57 | 22.61 |
| KOHI | Puaa | 8.42 | 1.85 | 28.96 | 30.64 |
| KOHI | 'Ulu | 9.45 | 2.00 | 36.82 | 27.42 |
| KULA | Breadnut | -- | -- | 38.94 | 25.96 |
| KULA | Fiti | 9.98 | 3.20 | 57.27 | 41.40 |
| KULA | Ma'afala | 10.07 | 3.24 | 58.44 | 37.92 |
| KULA | Otea | 8.77 | 3.42 | 49.46 | 32.28 |
| KULA | Puaa | 10.55 | 3.12 | 44.74 | 38.20 |
| KULA | 'Ulu | 9.77 | 3.20 | 51.31 | 34.80 |
| POAM | Breadnut | 7.09 | 3.09 | 43.50 | 28.53 |
| POAM | Fiti | 8.47 | 2.70 | 44.18 | 43.10 |
| POAM | Ma'afala | 8.32 | 2.62 | 46.84 | 32.55 |
| POAM | Otea | 7.82 | 2.98 | 53.34 | 38.25 |
| POAM | Puaa | 8.34 | 2.72 | 42.70 | 29.05 |
| POAM | 'Ulu | 8.68 | 2.82 | 51.69 | 50.33 |
